# The Relation Between White Matter Microstructure and Network Complexity: Implications for Processing Efficienc

**DOI:** 10.1101/299065

**Authors:** Ian M. McDonough, Jonathan T. Siegel

**Affiliations:** The University of Alabama, Department of Psychology, Tuscaloosa, AL 35487 USA; Center for Vital Longevity, School of Behavioral and Brain Sciences, University of Texas at Dallas, Dallas, TX 75235 USA

**Keywords:** diffusion tensor imaging, human connectome project, fMRI, multiscale entropy analysis, resting state networks, white matter microstructure

## Abstract

Brain structure has been proposed to facilitate as well as constrain functional interactions within brain networks. Simulation models suggest that integrity of white matter (WM) microstructure should be positively related to the complexity of BOLD signal—a measure of network interactions. Using 121 young adults from the Human Connectome Project, we empirically tested whether greater WM integrity would be associated with greater complexity of the BOLD signal during rest via multiscale entropy. Multiscale entropy measures the lack of predictability within a given time series across varying time scales, thus being able to estimate fluctuating signal dynamics within brain networks. Using multivariate analysis techniques (Partial Least Squares), we found that greater WM integrity was associated with greater network complexity at fast time scales, but less network complexity at slower time scales. These findings implicate two separate pathways through which WM integrity affects brain function in the prefrontal cortex—an executive-prefrontal pathway and a perceptuo-occipital pathway. In two additional samples, the main patterns of WM and network complexity were replicated. These findings support simulation models of WM integrity and network complexity and provide new insights into brain structure-function relationships.

## 1. Introduction

Some people process and analyze information quickly, whereas others do so more slowly. These individual differences in cognitive efficiency likely involve the coordination of many brain regions and are impacted by the integrity of both brain structure and function. Using the analogy of a road system, Nakagawa and colleagues [1] likened brain structure to street size (e.g., the amount of lanes) and brain function to traffic volume. They noted that street size both enables and limits the amount of traffic volume just as the integrity of brain structure enables and limits the amount of information flow between brain regions. They also emphasized that the effect of structure on function depends on the time scale with which traffic is measured—a property that has largely been ignored in human neuroscience. Consistent with this analogy, it is widely believed that brain structure impacts brain function. However, how structure impacts function and whether function impacts structure are far from well-understood.

One approach to understanding structure-function relationships has been to correlate the integrity of white matter (WM) microstructure and brain activity assessed in fMRI. WM integrity is most often measured using diffusion tensor imaging (DTI) to assess the degree of water diffusion within the white matter. For example, individuals with greater WM integrity (i.e., fractional anisotropy or FA) in frontoparietal regions exhibited increased brain activity in similar brain regions [2,3]. Few studies have investigated other measures of WM integrity such as axial diffusivity (AD), radial diffusivity (RD), or mean diffusivity (MD). Notably, one study did find that increased RD—representing greater demyelination—in lateral temporal cortex was associated with decreased brain activity in lateral frontal and cingulate cortices [4]. These findings suggest that individuals with greater WM integrity are able to recruit more neural resources and therefore might be able to process more information more quickly and efficiently than those with less WM integrity [5].

However, WM integrity in one region might also impact brain function in gray matter regions that are not directly connected to that region, but rather are more distal. That is, WM integrity might impact brain function in other functional networks. Studies have found when WM and brain activity have been assessed in non-adjacent regions, greater WM integrity has been associated with less brain activity [6–9]. This negative association might reflect a reduced reliance on or interactions between distal regions (i.e., more local processing), or could reflect a type of interference that prevents greater activity in distal regions within the same network or in outside networks (i.e., a disruption of distributed processing).

These studies illustrate the complicated relationship between brain structure and function and highlight the importance of separately characterizing local and distributed processing mechanisms. An emerging field within systems neuroscience has begun to investigate how temporal fluctuations within brain signals (i.e., predictability or lack thereof) is related to information processing. Specifically, researchers have proposed that less predictable fluctuations of brain activity are associated with richer [1,10–12] or more integrated information [13,14]. Unpredictable and highly complex biological systems often are related to a normal and healthy brain, whereas highly predictable or regular biological systems often are related to dysfunction and disease [15–17]. Critically, multiscale entropy (MSE) has been developed to estimate the complexity within temporal signals across multiple time scales [18]. MSE and its single scale counterpart, sample entropy (or SampEn), were originally used to understand temporal patterns of the human heartbeat [16,18]. Sample entropy estimates the irregularity of a time series, which increases as the amount of noise within a system increases. However, noise becomes quickly less complex towards coarser time scales because it has one single expected mean value [1,16]. Thus, the term “complexity” refers to the ability for a system to maintain irregularity across time series from fine to coarse. Over the last five years, researchers have extended the application of MSE and related entropy metrics to brain signals [e.g.,1,13,14,17,19,20,21]. This research has proposed that high frequency (fine) time scales and low-frequency (coarse) time scales represent local and distributed information processing, respectively [13,14,20]. Additionally, these measures of brain signal complexity offer complementary information that is not captured through other measures such as functional connectivity or standard deviation as evidenced by weak correlations between the various measures [19,21,22]. While many studies have used MSE to understand brain signal complexity using EEG [e.g., 13,14,17,19], only recently has research applied the same analysis tools to BOLD fMRI signals [e.g.,20,21,23].

The present study examined how WM integrity constrains brain function using MSE analyses, thus revealing its impact across multiple time scales. We base our predictions on simulations of WM integrity and complexity of the BOLD signal [1], but use non-simulated BOLD fMRI data to verify these simulations. According to Nakagawa and colleagues [1], people with greater WM integrity should have greater in neural complexity at fine time scales. They further suggested that these findings may emulate individual differences in brain and cognitive functioning (e.g., as with aging). For example, since WM integrity should be positively related to neural complexity, then as WM integrity declines with old age, brain function also would be less efficient, and ultimately lead to slower processing speeds—a general characteristic of aging [24,25]. Aging, however, is just one factor that can influence between-subject variability. Here, we tested the general notion that individual differences in WM integrity and neural complexity related to one another in a sample of healthy young adults from the Human Connectome Project (HCP, WU-Minn Consortium [26]). In accordance with these simulations, we predicted that individuals with greater WM integrity would exhibit *greater* local network complexity in the HCP data. However, because the data used in the present study can afford a larger range of time scales than that used in the model simulations, we tested a new hypothesis that individuals with greater WM integrity would show *reduced* distributed network complexity than individuals with less WM integrity. Our rationale for this reverse effect stems from previous research revealing a) an inverse relationship with WM integrity and distal brain activity [5] and b) an inverse relationship between functional connectivity and neural complexity at fine (local) and coarse (distributed) time scales [20]. To understand these network dynamics, multivariate network analyses were implemented. Lastly, we tested our predictions in three samples to generalize our results. These finding should shed light on the impact of WM integrity on brain function and help further our understanding of patterns of MSE in the BOLD signal.

## 2. Materials and Methods

### 2.1 Participants

Data was taken from participants in the HCP, which is a long-term study to explore human brain circuits. Participants were young adults that were relatively healthy and free of a prior history of significant psychiatric or neurological illnesses, but could have a history of smoking, heavy drinking, or recreational drug use. All subjects gave their written informed consent for inclusion before they participated in the study. The study was conducted in accordance with the Declaration of Helsinki, and the protocol was approved by the Ethics Committee of Washington University, St. Louis and The University of Alabama.

Four hundred and ninety adults, some of which were siblings, were included in the study if they had available DTI and fMRI data. From this total, three subgroups were formed (see the sample characteristics for the final samples in Table 1). The first group consisted of 123 unrelated individuals and was used for the primary analyses. The second and third groups were used for replication purposes. Due to the nature of the HCP data, these other two groups consisted of related individuals (both related to each other within the subgroup as well as related to the first group). Of these two groups, we were able to match the second group to the first on demographic factors including age, education, sex, and racial status (white or non-white). The third group was older than the other two groups, had more education, and was more likely to be white (all *p*’s < .001). There were no differences in sex between the last group and the first two groups (all *p*’s > .17).

**Table 1.** Sample Characteristics Across Three Samples.

|  | Primary | Matched | Non-Matched |
| --- | --- | --- | --- |
| N | 121 | 122 | 121 |
| Age (SD) | 27.97 (3.14) | 28.11 (3.54) | 30.37 (3.04) |
| Years of Education (SD) | 14.38 (2.28) | 14.30 (2.26) | 15.69 (1.52) |
| Sex (% Female, % Male) | 57%/43% | 63%/37% | 65%/35% |
| Race |  |  |  |
| White (%) | 66% | 68% | 92% |
| Black (%) | 24% | 30% | 6% |
| Asian/Pacific Islander (%) | 3% | 1% | 2% |
| Other (%) | 7% | 1% | 1% |

Note. SD = standard deviation.

### 2.2 DTI procedures

Diffusion data were collected using a single-shot, single refocusing spin-echo, echo-planar imaging sequence with 1.25 mm isotropic spatial resolution (TE/TR = 89.5/5520 ms, FOV = 210 × 180 mm). Three gradient tables of 90 diffusion-weighted directions and six b = 0 images each were collected with right-to-left and left-to-right phase encoding polarities for each of the three diffusion weightings (b = 1000, 2000, and 3000 s/mm^2^). HCP preprocessed DTI images were used which included correcting for B_0_, susceptibility artifact, and eddy current distortions. For a more detailed description of the DTI acquisition and preprocessing procedures [27,28]. Linearly fitting a diffusion tensor model to the DTI images resulted in FA, eigenvector, and eigenvalue maps that were then used to estimate axial diffusivity (AD; λ_1_), radial diffusivity (RD; (λ_2_ + λ_3_)/2)), and mean diffusivity (MD; (λ_1_ + λ_2_ + λ_3_)/3). Using Tract Based Spatial Statistics (TBSS) [29], FA images were skeletonized and normalized to the FMRIB58 template using a non-linear image registration tool (FNIRT). Using the mean FA image, a skeletonized mask was created representing the tract centers common to all participants. Mean DTI values for each measure were then extracted from regions of interest based on the JHU atlas. The tracts included 30 major association, projection, and callosal white matter tracts, which were then extracted using masks from the JHU atlas. Due to computational demands, we had to exclude some tracts. We chose to first exclude tracts that had few voxels (thus potentially leading to poor estimations) or were not immediately relevant to the higher order cognitive networks of interest (exclude tracts included Fornix (cres)/Stria Terminalis, Cerebellar tracts).

### 2.3 fMRI procedures

All data were acquired on a Siemens Skyra 3T scanner housed at Washington University in St. Louis. The scanner had a customized SC72 gradient insert and a customized body transmitter coil with 56 cm bore size (diffusion: Gmax = 100 mT/m, max slew rate = 91 mT/m/ms; readout/imaging: Gmax = 42 mT/m, max slew rate = 200 mT/m/ms). The HCP Skyra had the standard set of Siemen’s shim coils (up to 2nd order) and used Siemen’s standard 32 channel head coil. BOLD fMRI data were acquired using a T2*-weighted gradient-echo EPI sequence with 72 axial slices per volume, 104 × 90 matrix (2.0×2.0×2.0 mm^3^), FOV=208 mm, TE=33.1 ms, TR=720 ms, FA=52°. Across four scanning sessions of 15 minutes each, a total of 4800 frames were acquired. Participants were instructed to keep their eyes open and focused on a bright cross-hair on a dark background. Across sessions, oblique axial acquisitions alternated between phase encoding in a right-to-left direction and phase encoding in a left-to-right direction.

Postprocessed fMRI datasets were used in the present study, which consisted of standard processing methods using FSL [30,31]. Below briefly summarizes the HCP processing pipeline [32]. First, gradient-nonlinearity-induced distortion was corrected for all images. Next, FMRIB’s Linear Image Registration Tool (FLIRT) was used for motion correction using the single-band reference (SBRef) image as the target. The FSL toolbox “topup” [33] was used to estimate the distortion field in the functional images. The SBRef image was used for EPI distortion correction and is registered to the T1w image. One-step spline resampling from the original EPI frames to MNI space was applied to all transforms. Lastly, image intensity was normalized to mean of 10,000 and bias field was removed. Data was cleaned using ICA+FIX [34,35], which included linear detrending, regression of 24 motion parameters, and ICA noise components removed. This method better removes artifacts than regressing out white matter and/or CSF signal directly, as well as using the “scrubbing” method [36]. Global signal was not removed. Importantly, while choosing methods to preprocess resting-state fMRI data can be controversial, studies have shown that group differences in MSE are quite robust to variations in preprocessing methodologies [37].

### 2.4 Multiscale entropy (MSE) analysis

The processed time series were extracted at each point on the normalized brains and averaged together if they fell within distinct resting-state networks (RSN) using the parcellation of 24 networks from the Power atlas [38]. This network approach was taken since networks are often defined by the similarity of their time series, thus resulting in a similar pattern of complexity within a given network [20]. Deriving MSE from networks offers a simplified and theoretically driven approach to investigating MSE across the brain. After the time series from each RSN was extracted, the complexity of each network (separately for each of the four resting-state scans) was estimated using MSE. MSE estimates sample entropy at different time scales. First, fine to more coarse-grained time series were created by down-sampling the original time series (i.e., averaging neighboring data points within non-overlapping windows). Second, sample entropy was estimated for the time series at each time scale (1 to 25 scales). Sample entropy is defined as the natural logarithm of the conditional probability that a given pattern of data of a specified length (*m*) repeats at the next time point for the entire time series at a given scale factor (of a dataset with a total length *N*). It considers subsequent patterns to be a repeat of the given pattern if they match within a certain tolerance (*r*) such that larger tolerance values increase the number of matches [39,40]. A time series with a greater number of pattern matches is more predictable and the entropy value is lower (less complexity). In contrast, a smaller number of pattern matches is characterized as being less predictable, yielding a greater entropy value (greater complexity). We selected our parameters based on those used in prior studies investigating MSE using fMRI, *m* = 2 and *r* = .5 [20, 41,42]. Previous investigations using the data from the HCP revealed qualitatively similar patterns of MSE estimations regardless of the parameters chosen [20]. This consistency is similar to other studies who have found that the choice of parameters leads to relatively robust estimations over a broad range of possible values [39,40,42]. The resulting MSE estimations for each of the four runs were then averaged together to obtain a more robust estimation of MSE for each time scale within each network.

### 2.5 Partial least squares (PLS) analyses

PLS was used to determine the regional WM associations with MSE across RSNs. PLS is a multivariate technique designed to identify latent factors that account for most of the variance in a data set [43]. For the PLS analysis, the X matrix was organized in the form of [Subjects x Time Scale in Network], resulting in a 123 × 600 matrix that represented network complexity in the 24 RSNs with 25 time scales for each network for each subject. The Y matrix was organized in the form of [Subjects x tracts in DTI measure], resulting in a 123 × 120 matrix that represented DTI values across the 30 different tracts for FA, AD, MD, and RD for each subject. Each matrix was grand-mean centered and normalized prior to conducting the analysis.

The cross product of the X and Y matrices was then decomposed into a set of mutually orthogonal factors using singular value decomposition, resulting in a set of orthogonal latent variables (LVs). An LV consists of three components: 1) a singular value, 2) a vector of weights representing the pattern of time scales in the LV (i.e., salience values), and 3) a vector of weights representing the degree to which each subject expresses the given LV (i.e., brain scores). Brain scores were calculated by multiplying the salience scores by the network complexity values for each subject and then summing these values. Each LV was statistically evaluated two ways. First, we assessed the significance of the relationship between network complexity and the DTI measures by computing 10,000 permutation tests in which network complexity values were randomly assigned within subjects. A measure of significance was calculated by estimating the proportion of times the permuted singular value was higher than the observed singular value. Second, to assess the reliability of the corresponding distribution across subjects (i.e., saliences), we resampled subjects with replacement (10,000 bootstrap samples). A bootstrap ratio was then calculated by dividing the saliences by the standard error of the generated bootstrap distribution. The bootstrap ratio is approximately equivalent to a z-score, whereby an absolute bootstrap ratio greater than 1.96 corresponds roughly to *p* < .05. When bootstrapped confidence intervals did not cross zero, we considered those values to be statistically significant.

## 3. Results

After forming the three groups, preliminary analyses across all three groups revealed five outliers that had at least one DTI measure (averaged across the whole brain) greater than 7.5 standard deviations from the mean, resulting in final group sizes of 121 (primary group), 122 (matched group), and 121 (non-matched group). We first report the PLS analyses on the primary group to determine the extent that network complexity was associated with WM integrity. Finally, we report whether any of the findings replicated in the separate matched and non-matched samples.

### 3.1 Network complexity and WM integrity

The PLS analysis resulted in four significant LV’s that explained 78.57%, 10.01%, 5.27%, and 2.33% of the covariance in the data, respectively. Because the first two LV’s comprised almost 90% of the covariance, we focused on these LV’s. For the first LV (*p* = .009), a Pearson correlation conducted between the network-complexity brain scores and the WM brain scores confirmed a moderate effect size between the two factors (*r* = .39, *p* < .001). As seen in Figure 1 (top panel), the first LV pattern was characterized by greater network complexity at fine time scales and lower network complexity at mid and coarse time scales. Furthermore, this pattern was strongest in the following networks: default mode, frontoparietal, salience, cingulo-opercular, dorsal and ventral attention, “unknown Nelson 2010”, and “unknown with memory retrieval”. These functional networks have been implicated in higher-order cognition [44–47] and most of these networks contain key regions within the prefrontal cortex. In regards to WM, this LV pattern was associated with lower FA and lower AD across most of the tracts, and lower MD in a smaller subset of tracts. In contrast, this LV was associated with weaker and more mixed relationships with RD. The tracts that most exhibited this WM pattern (and had confidence intervals that did not include zero) consisted of projection and association tracts and were left lateralized including left anterior corona radiata, posterior corona radiata, posterior thalamic radiata, external capsule, superior fronto-occipito fasciculus, superior longitudinal fasciculus, uncinate fasciculus. The right hemisphere consisted of anterior corona radiata, and superior longitudinal fasciculus. Like network complexity, many of these WM tracts contain connections with prefrontal cortex. In sum, individuals with greater network complexity at fine time scales and lower complexity at mid and coarse time scales also had lower FA, AD, and MD values.

**Figure 1.**
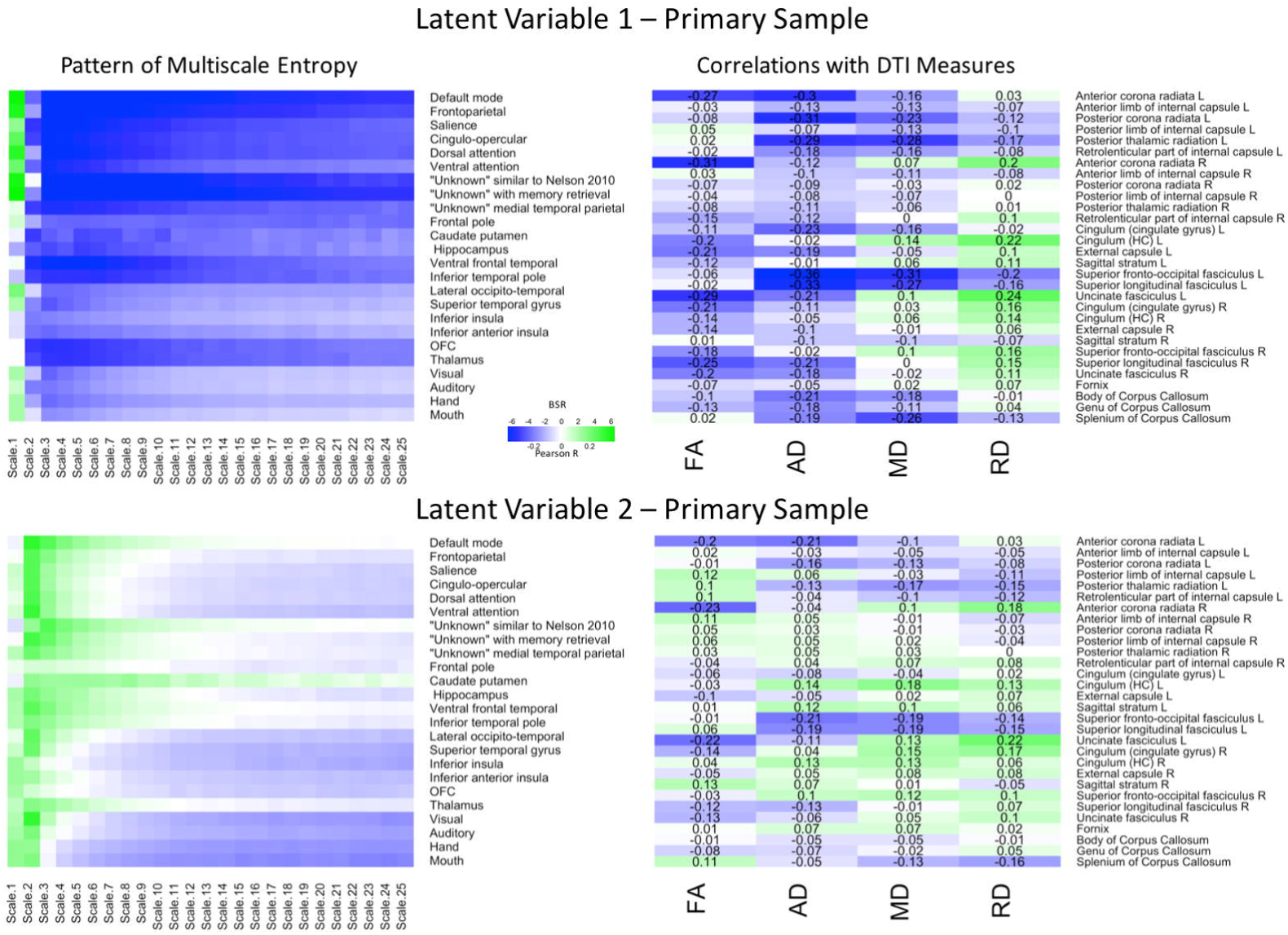
LV 1 (top) and LV 2 (bottom) from the partial least squares analysis in the primary sample. The left panels represents bootstrap ratio (BSR) values for network complexity as measured by multiscale entropy across 25 time scales. The right panels represents the correlation values for each of the DTI measures (FA, AD, MD, and RD). For the LV 1, greater network complexity values at the fine time scale (shown in green in the left panel) were associated with lower FA, AD, and MD values (shown in blue in the right panel), but higher RD values in many of the tracts (shown in green in the right panel). In contrast, greater network complexity values at the mid and coarse time scales (shown in blue in the left panel) were associated with lower FA, AD, and MD values, but higher RD values. For LV 2, greater network complexity values at the fine time scales, but lower network complexity at mid and coarse time scales were associated with both lower (in blue) and higher (in green) DTI values depending on the measure and tract. LV 1 = latent variable 1; LV 2 = latent variable 2; FA = fractional anisotropy; AD = axial diffusivity; MD = mean diffusivity; RD = radial diffusivity.

The second LV (*p* = .019) had a weak effect size between between network complexity and white matter (*r* = .25, *p* < .005). As shown in Figure 1 (bottom panel), this LV pattern was characterized by greater network complexity at the second time scale (across almost every network) and lower network complexity at coarse time scales largely in sensorimotor networks including visual, auditory, hand, and mouth networks. The pattern of white matter was characterized by lower FA, AD, and MD, but greater RD, in only a few major tracts. Specifically, the FA relationships (that had confidence intervals excluding zero) were lowest in left and right anterior corona radiata, and left uncinate fasciculus. The AD and MD relationships were lowest in left anterior and posterior corona radiata, left superior fronto-occipital fasciculus, and left superior longitudinal fasciculus. The RD relationships were greatest (and negative) in the right anterior corona radiata, left uncinate fasciculus, and right cingulum. Together, the results from these two LV’s are consistent with the predictions that greater WM integrity would be associated with greater local network complexity (i.e., entropy at fine time scales), but less distributed network complexity (i.e., entropy at coarse time scales).

### 3.2 Replication of effects: Matched sample

We next conducted the same PLS analysis on the matched sample (Figure 2). The first LV explained 66.55% of the covariance and was marginally significant (*p* = .057). Importantly, a Pearson correlation between the network-complexity brain scores and the WM brain scores exhibited a similar, moderately sized correlation (*r* = .31, *p* < .001). The pattern of network complexity also was quite similar as in the primary sample; major higher-order networks involving prefrontal cortex were characterized by greater network complexity at the finest time scale and lower network complexity at the mid and coarse time scales. Like in the primary data set, this LV also was characterized by lower AD and MD in many of the same tracts including left anterior left anterior corona radiata, posterior thalamic radiata, superior fronto-occipito fasciculus, and superior longitudinal fasciculus. Unlike the primary group, the first LV in the matching group was characterized by higher FA values and many more consistent lower RD values. Thus, while differences did exist, the network-complexity measures and two of the WM measures (AD and MD) were quite consistent with the analysis in the primary group.

**Figure 2.**
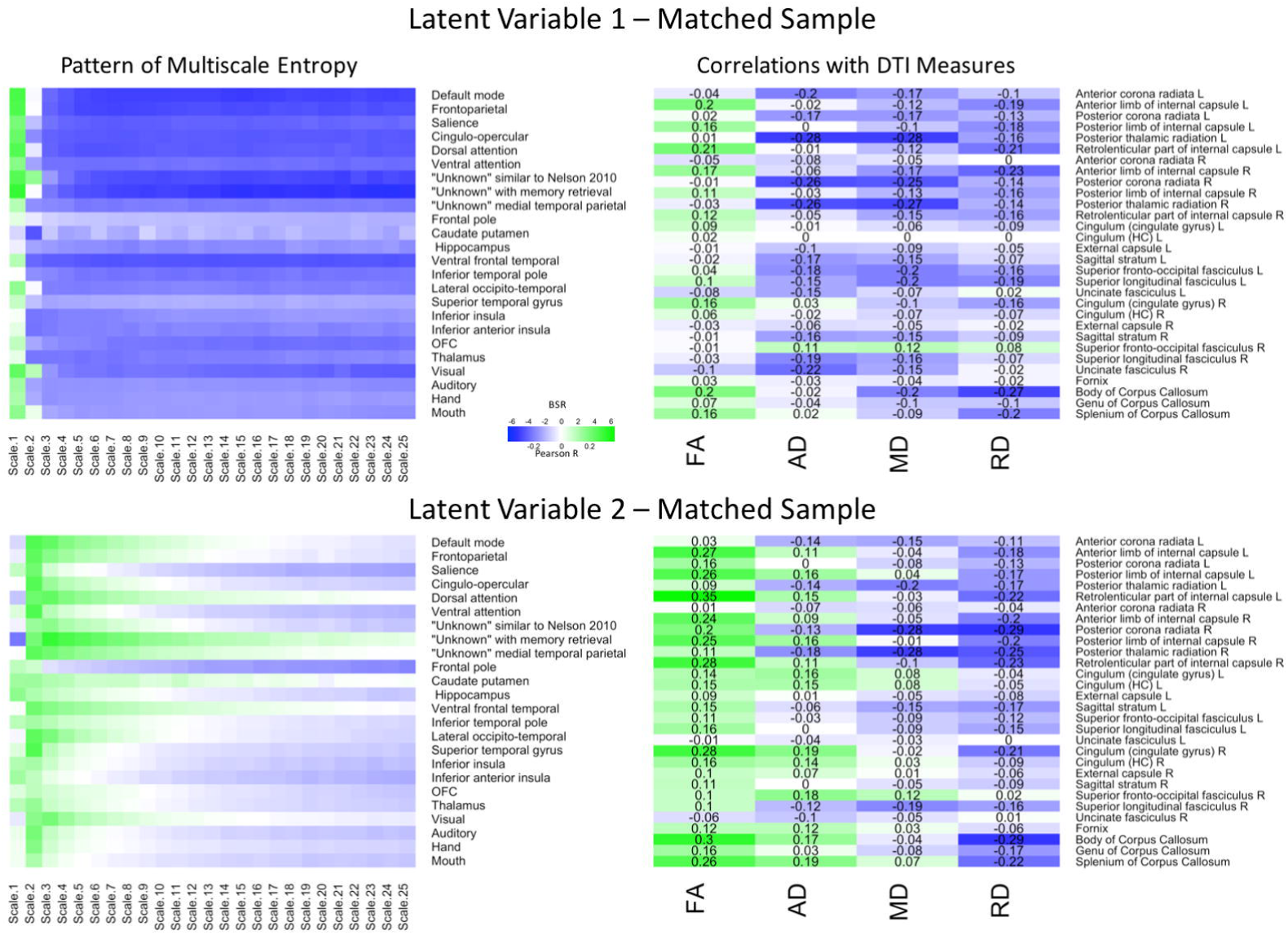
LV 1 (top) and LV 2 (bottom) from the partial least squares analysis in the matched sample. The left panels represents bootstrap ratio (BSR) values for network complexity as measured by multiscale entropy across 25 time scales. The right panels represents the correlation values for each of the DTI measures (FA, AD, MD, and RD). For the LV 1, greater network complexity values at the fine time scale (shown in green in the left panel) were associated with lower AD, MD, and RD values (shown in blue in the right panel), but higher FA values in many of the tracts (shown in green in the right panel). In contrast, greater network complexity values at the mid and coarse time scales (shown in blue in the left panel) were associated with lower AD, MD, and RD values, but higher FA values. For LV 2, greater network complexity values at the fine time scales, but lower network complexity at mid and coarse time scales were associated with lower MD and RD values (in blue), but higher FA and AD values (in green) across many of the tracts. LV 1 = latent variable 1; LV 2 = latent variable 2; FA = fractional anisotropy; AD = axial diffusivity; MD = mean diffusivity; RD = radial diffusivity.

The second LV explained 21.89% of the covariance (*p* < .001). The Pearson correlation between the brain scores revealed a weak correlation as in the second LV from the primary group (*r* = .26, *p* < .005), and the pattern of network complexity also was similar. Network complexity was characterized by greater complexity at the second time scale across most networks and lower complexity at mid and coarse time scales in many of the networks, but failed to reach significance in the sensorimotor networks (BSR ≤ 1.96). However, the pattern of WM was different. Specifically, this pattern consisted of greater FA and AD values, and lower MD and RD values. Only the MD values were in a direction consistent with the primary group, but the tracts were different (largest correlations in right posterior corona radiata and right posterior thalamic radiation). Thus, while the second LV did show a significant relationship between network complexity and WM integrity, the different white matter tracts support only a partial replication.

### 3.3 Replication of effects: Non-matched sample

Finally, we conducted the same sets of analyses on the non-matched sample (Figure 3). The first LV (*p* = .005) explained 77.85% of the covariance. The Pearson correlation between two brain scores had a moderate effect size (*r* = .37, *p* < .001). The network-complexity patterns largely resembled the previous two analyses, whereas the WM patterns largely resembled the previous, matched-sample analysis. Specifically, the pattern consisted of greater FA, but lower AD, MD, and RD values in many of the same tracts as the matched-sample analysis. The second LV (p < .001) explained 13.98% of the covariance and the Pearson correlation between the two brain scores had a weak effect size (*r* = .23, *p* = .011). The pattern of network complexity was similar as the other two analyses with the second time scale across most networks, but the mid and coarse time scales in the sensorimotor networks failed to reach significance (as in the Match-sample analysis). The pattern of white matter consisted of greater FA, mixed AD, and lower RD and MD values. In relation to the other analyses, the specific tracts that show negative relationships with AD and MD values are consistent with the analyses in the primary group and the tracts with the greatest FA values were consistent with the analysis in the matched sample.

**Figure 3.**
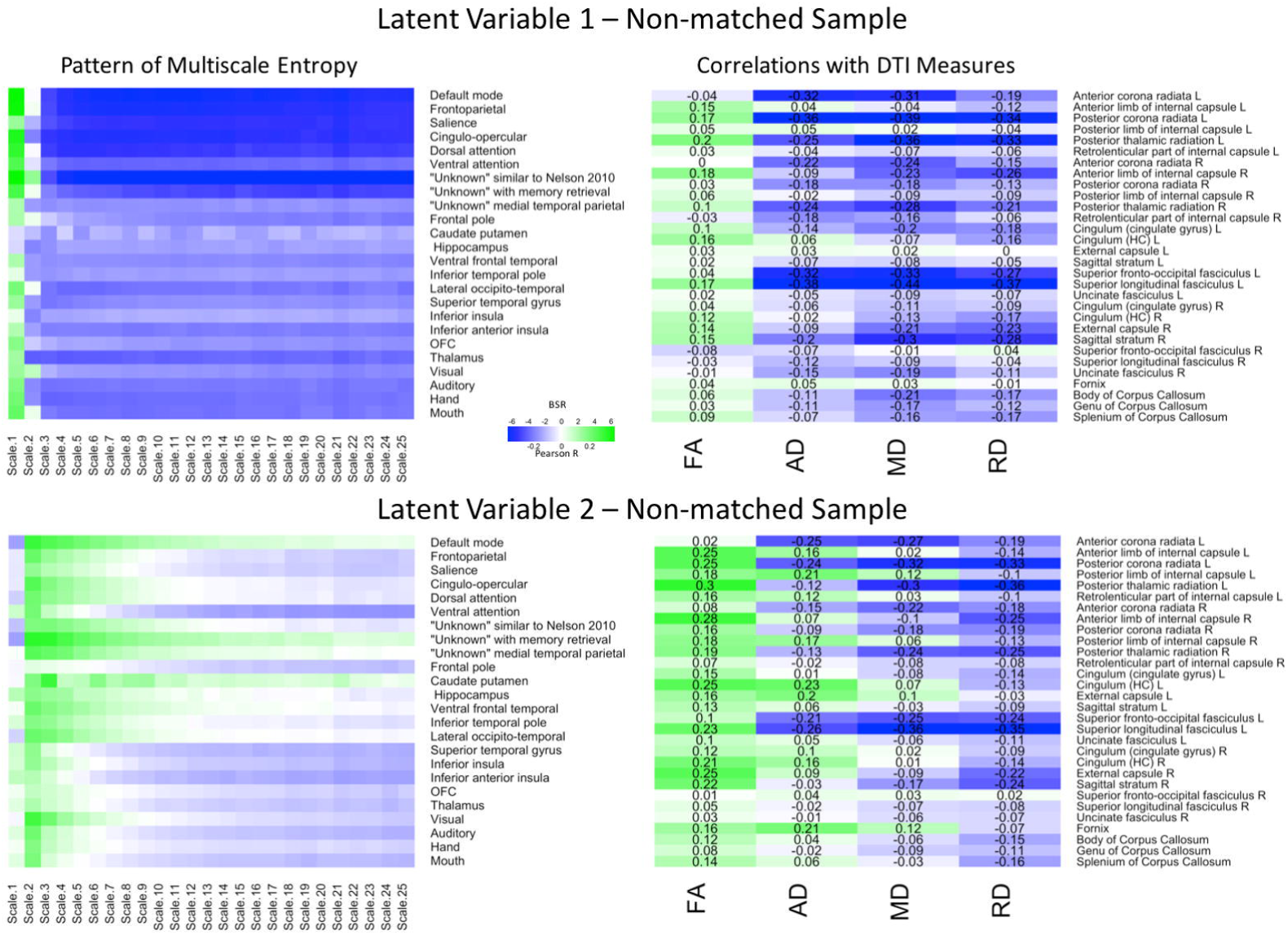
LV 1 (top) and LV 2 (bottom) from the partial least squares analysis in the non-matched sample. The left panels represents bootstrap ratio (BSR) values for network complexity as measured by multiscale entropy across 25 time scales. The right panels represents the correlation values for each of the DTI measures (FA, AD, MD, and RD). For the LV 1, greater network complexity values at the fine time scale (shown in green in the left panel) were associated with lower AD, MD, and RD values (shown in blue in the right panel), but higher FA values in many of the tracts (shown in green in the right panel). In contrast, greater network complexity values at the mid and coarse time scales (shown in blue in the left panel) were associated with lower AD, MD, and RD values, but higher FA values. For LV 2, greater network complexity values at the fine time scales, but lower network complexity at mid and coarse time scales were associated with lower MD and RD values (in blue), but higher FA values (in green) across many of the tracts. AD measures showed both positive and negative associations with network complexity in LV2. LV 1 = latent variable 1; LV 2 = latent variable 2; FA = fractional anisotropy; AD = axial diffusivity; MD = mean diffusivity; RD = radial diffusivity.

### 3.4 Cross-Sample Summary

To better depict which relationships were significant across each of the three samples, conjunction maps were created for both LV’s (Figure 4). For network complexity, BSR’s values at least |1.96| (corresponding to *p* = .05) that were consistent across two or three samples were included. For DTI measures, *r* values of at least |.20| (also corresponding to *p* = .05) that were consistent across two or three samples were included. For LV 1, the network complexity pattern was consistently significant in all three samples for almost every time scale and most networks with the largest exception being in the sensory networks (visual, auditory, hand, mouth). However, the DTI pattern was only consistent across the three samples in left anterior corona radiata (AD), left posterior thalamic radiation (AD and MD), left superior fronto-occipital fasciculus (MD), left superior longitudinal fasciculus. For LV 2, network complexity was replicated in time scale 2 for the majority of the networks (many of which overlapped with the significant networks in LV 1), followed by time scale 3 and 4. None of the DTI measures for LV 2 were consistently significant across the three samples, but several were consistent in two of the three samples. These regions included left anterior corona radiata (AD), left and right posterior thalamic radiata (MD), and left superior frontal-occipital fasciculus (AD). However, it should be noted that this conjunction analysis is quite conservative, equating to a joint *p*-value of .000125 for commonalities across all three samples and a joint *p*-value of .0025 for commonalities across two of the three samples. These stricter thresholds explain the sparse overlap across the three samples.

**Figure 4.**
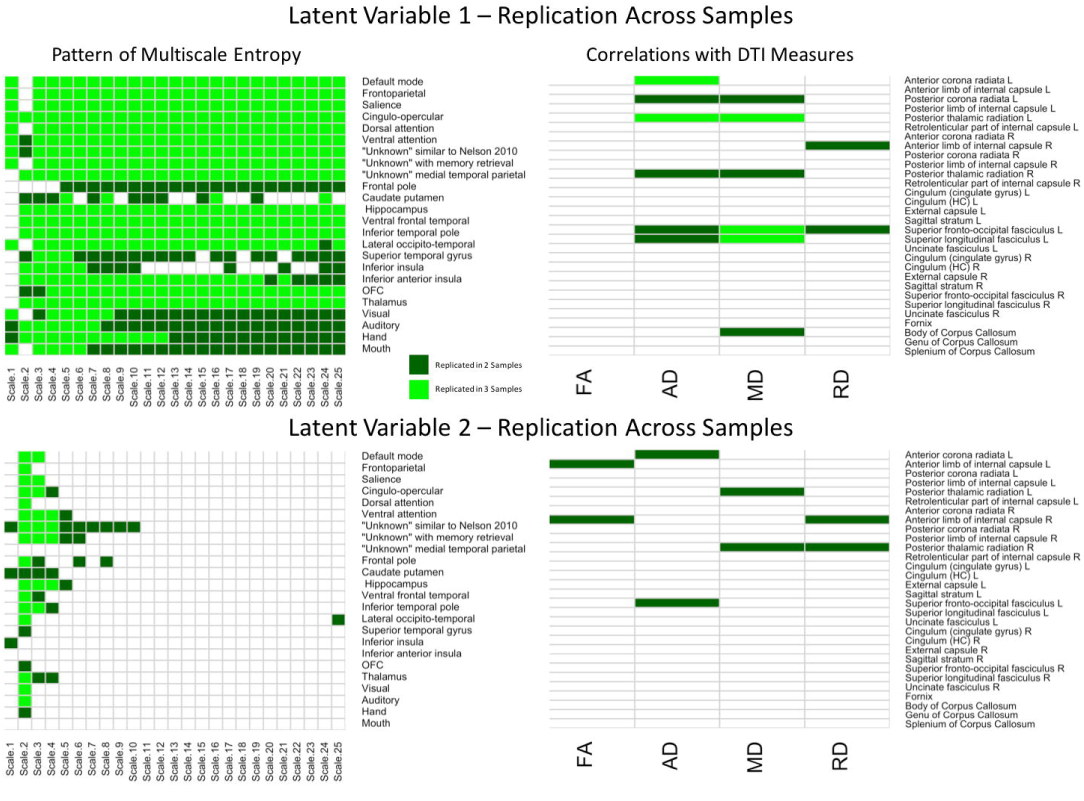
Cells in green represent replications of significant correlations between MSE and WM microstructure across all three samples for LV 1 (top) and LV 2 (bottom). The left panels represent brain networks across 25 time scales. The right panels represent the DTI measures (FA, AD, MD, and RD). LV 1 = latent variable 1; LV 2 = latent variable 2; FA = fractional anisotropy; AD = axial diffusivity; MD = mean diffusivity; RD = radial diffusivity.

## 4. Discussion

While it is clear that brain structure influences brain function, how this happens is far from understood. In the present study, we took an individual differences approach to test the notion that people with greater WM integrity would exhibit greater localized processing, but less distributed processing than people with less WM integrity. To do this, we assessed brain structure using four measures of WM integrity including FA, AD, MD, and RD—all of which provide complementary information regarding the integrity of WM. To capture brain function across multiple time scales, we used multiscale entropy to estimate the complexity of the BOLD signal across 24 resting-state networks. This network complexity might represent the richness or integration of information processing signals that underlie processing efficiency [1,13,14,20]. Using a multivariate analysis technique, we found that WM integrity across many tracts was significantly associated with network complexity across many resting-state networks (RSNs). We elaborate on the details of these findings below.

### 4.1 Latent Variable 1: White Matter Integrity Influences Network Complexity in the Prefrontal Cortex Networks

We first aimed to better understand how brain structure integrity influences the capacity of information processing within functional brain networks via network complexity measures. In the first latent variable of the PLS analysis, greater network complexity at fine time scales and lower network complexity at mid and coarse time scales were associated with greater WM integrity (i.e., lower AD and MD)—greater axonal density or fiber coherence [48–50]. This latent variable was consistent across the three samples. Better WM integrity might facilitate interconnectivity among local neural populations, perhaps at the expense of long-range interactions across distributed neural populations [13,14,20]. This idea is consistent with previously found positive correlations between WM integrity and brain activity in regions spatially adjacent to one another, but negative correlations when the regions being measured are more distal from one another [5].

The WM tracts that had the greatest associations with network complexity can be divided into two categories: anterior association tracts and posterior projection tracts. The anterior association tracts—connected to the prefrontal cortex [51,52]—consisted of the left superior fronto-occipital fasciculus and the superior longitudinal fasciculus and are related to processes including working memory [53,54]. The posterior projection tracts comprise of portions of the internal capsule (anterior corona radiata and posterior thalamic radiation), connecting the thalamus to both prefrontal and occipital cortex. These tracts have been primarily implicated in perceptual and motor functions [3]. In regards to brain function, the latent variable indicated that the strongest effects were in RSNs that included the prefrontal cortex including the default mode, frontoparietal, salience, cingulo-opercular, dorsal attention networks. While there are many regions that comprise each of these networks, we focus on the prefrontal cortex which is shared by all of the networks. Moreover, each of these RSNs has been implicated in higher-order cognition (e.g., attention, working memory, executive function), which often involves the use of the prefrontal cortex [44–47]. In regards to WM integrity, greater WM integrity fosters faster information processing. For example, lower values of AD and MD in various tracts including the anterior thalamic radiation have been associated with faster response times during a speeded continuous performance task [55]. Other studies also have found relationships with WM integrity (i.e., FA) and processing speed [56,57] and attention [58,59]. Importantly, the tracts in these studies overlapped with the tracts here including the superior longitudinal fasciculus and corona radiata. Because of their broad nature, these structural and functional relationships may form the basis for processing efficiency across a variety of cognitive domains. Together, these results suggest that two primary components of brain structure—an executive-prefrontal component and a perceptuo-occipital component—affect brain function within the prefrontal cortex (see Figure 5).

**Figure 5.**
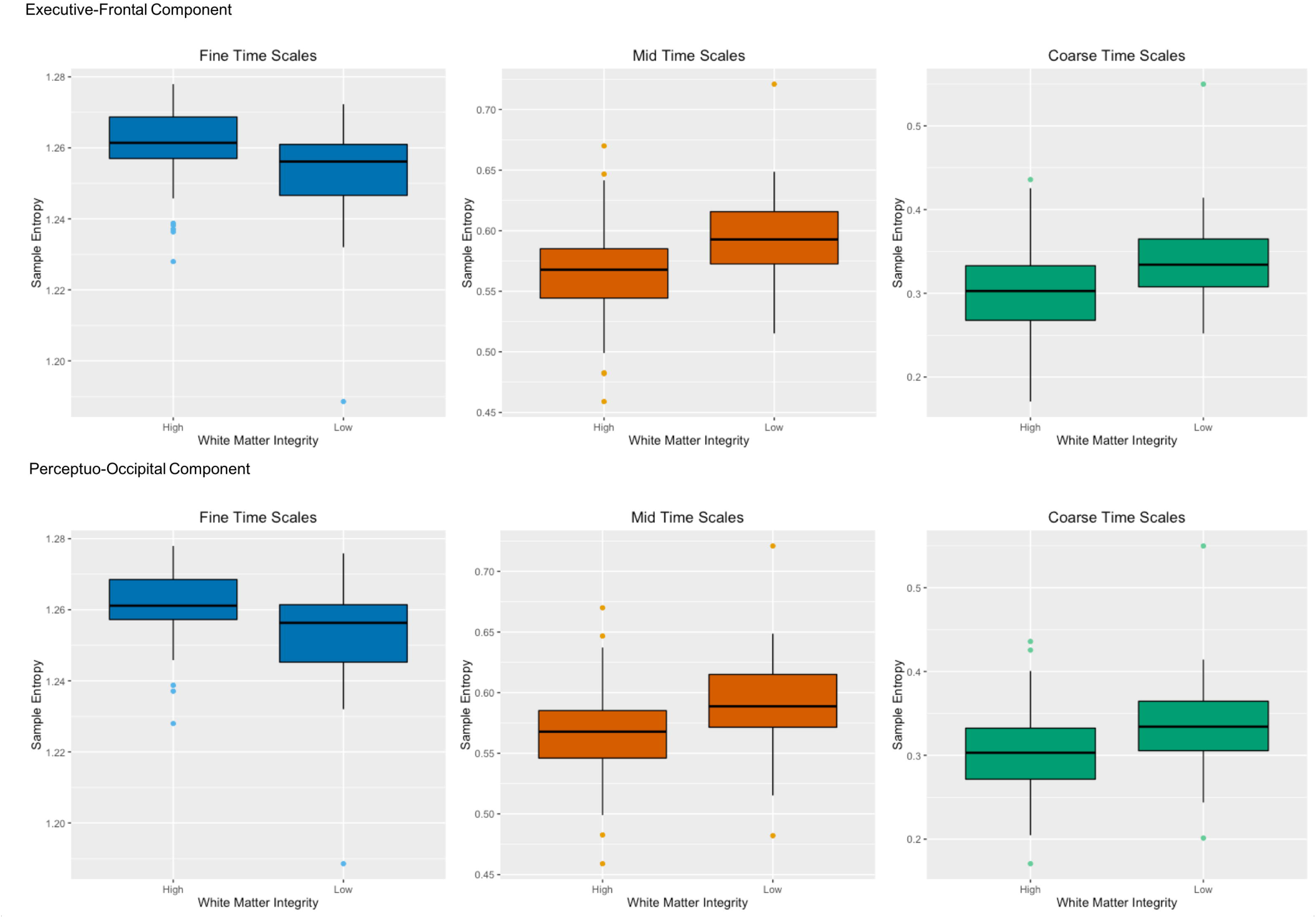
Two primary components of brain structure—an executive-prefrontal component and a perceptuo-occipital component—affect network complexity within the prefrontal cortex. The top panel shows high and low white matter integrity from anterior white matter tracts (left superior longitudinal fasciculus and left superior fronto-occipital fasciculus) for both AD and MD on the x-axis with sample entropy on the y-axis for fine time scales (scale 1), mid time scales (scales 3-14) and coarse time scales (scales 15-25). The bottom panel shows the same for high and low white matter integrity from posterior white matter tracts (left anterior corona radiata and left posterior thalamic radiation) for both AD and MD. Greater white matter integrity was associated with higher network complexity than poorer white matter integrity at fine time scales, but was associated with lower network complexity at mid and coarse time scales.

### 4.2 Latent Variable 2: White Matter Integrity Affects Complexity Independent of Network

After accounting for the majority of the covariance between brain structure and function, a second latent variable revealed that greater WM integrity (i.e., lower AD and MD) was associated with greater network complexity at fine scales, although the exact time scale was shifted to the second time scale. While many WM tracts were associated with this latent variable pattern, the left superior fronto-occipital fasciculus and superior longitudinal fasciculus were most consistently found across the three samples. Both of these tracts contain long-range fibers and connect the prefrontal cortex with other cortical brain areas including parietal and occipital regions [51,52,60]. Neural complexity associated with this latent variable was found across almost every network and the three samples in the second time scale.

### 4.3 Interpreting Network Complexity

Previous studies suggest that greater local WM integrity is associated with greater local brain activity, but reduced distal brain activity [5]. While the present results are consistent with those notions, the network complexity measures used here provide a complementary approach to measure local and distal interactions between brain regions and provide additional information as to how the richness of information processing is impacted by WM integrity. Also novel is the interaction between time course and the direction of the structure-function relationship—an effect that would not have been easily predicted directly from previous studies using non-complexity measures. As reviewed by Shen et al. [61], some studies have suggested that brain function emerges from brain structure, but is best reflected at coarse timescales and is only weakly correlated at finer time scales [62]. The present findings suggest that brain structure at multiple time scales including finer time scales as applicable to the BOLD signal, which might be considered a coarser time scale in itself. Specifically, we found that greater complexity at fine time scales is associated with greater within-region processing whereas greater complexity at coarser time scales is associated with more distributed processing across regions or networks [13,14,20]. For example, McIntosh and colleagues [13] found that complexity at fine time scales was associated with more within-hemisphere (local) functional connectivity, but complexity at coarse time scales was associated with more between-hemisphere (distributed) functional connectivity. Thus, the greater WM integrity would be interpreted as facilitating local interconnectivity and inhibiting or interfering with distributed connectivity. This interpretation assumes that individual differences in WM integrity lead to *quantitative* differences functional interactions. That is, the brain operates in fundamentally similar ways whether one has a high or low level of structural integrity.

A different interpretation is that people with high or low level of structural integrity utilize functional networks in *qualitatively* different manners. People with greater WM integrity might rely more on local neural processing, whereas people with poorer WM integrity might rely more on distributed neural processing. To the extent that local WM integrity is high, less information might be lost when “in transit” from one neighboring gray matter region to the next, resulting in a maintenance of rich information processing. In contrast, if local WM integrity is low, more information might be lost when it is being processed within neighboring regions, resulting in less rich information processing. In response to this degraded information, people with poor WM integrity might need to draw resources from other brain regions in surrounding areas (non-local).

Relying less on local neural processing and more on distributed neural processing has implications for individual differences in intelligence and network resiliency to aging or disorders that impact that brain. For instance, studies have shown that more intelligent young adults show less distributed neural processing [63,64,65]. However, as people get older and their intellectual faculties begin to decline, older adults rely on more distributed neural processing, which has been argued to either compensate for structural declines [66,67] or simply be a measure of less efficiency in the neural system [68,69]. Consistent with these ideas, Daselaar et al. [70] recently found that older adults with lower executive function exhibited an increase in distributed neural processing, which was in turn associated with reduced white-matter integrity. Gao et al. [71] found similar results in older adults and patients with Alzheimer’s disease. They found that white-matter integrity of short-range fibers contributed to more brain activity and lower cognitive efficiency.

A limitation to these interpretations, however, is the fact that much of the foundational work investigating the nature of MSE has been done in EEG and MEG, leaving much research to be conducted on the exact nature of MSE patterns in the BOLD signal [20,22].

### 4.4 Reliability and Strength of the Findings

Neuroscience and psychological research has been under fire for failing to replicate key findings [72,73]. Here, we used multiple methods to obtain reliable results including using bootstrapping, large sample sizes (relative to most neuroimaging studies), and three different samples. With stringent criteria for replication, we found four of seven of the strongest correlations were replicated twice and three of seven of the strongest correlations were replicated once. Specifically, in Figure 1 we can see that the strongest correlations were for AD in left superior fronto-occipital fasciculus (r = -.36), left superior longitudinal fasciculus (r=-.33), left anterior corona radiata (r = -.30), left posterior corona radiata (r = -.31), and left posterior thalamic radiation (r = -.29), and for MD in left superior fronto-occipital fasciculus (r = -.31) and left superior longitudinal fasciculus (r=-.27). Given that the first seven of the strongest correlations were replicated at least once at a very stringent threshold, we feel that reproducibility was quite good for the strongest relationships.

Despite these strengths, we only partially replicated our analyses. While the association between network complexity and two of the measures of WM integrity (AD and MD) replicated across the three samples, the direction of FA and RD were more similar between the two replication samples than in the primary sample. Because of this inconsistency, we have chosen not to interpret the FA and RD effects. It may be the case that some measures are not as reliable as one might hope and many studies that do not attempt to take these steps to enhance or validate its reliability may lead to inaccurate inferences. It is also possible that critical characteristics (other than age, education, sex, and race) that differed across the three samples masked potential relationships.

The strength of even the most reliable relationships was weak to moderate. On the one hand, these low correlation strengths might indicate that WM integrity *does not* influence network complexity (i.e., there is not a one-to-one connection between the two modalities). Instead, other factors might have a stronger impact on network complexity including the density of neurons, the number of synapses per neuron, blood flow, brain metabolism, among others [74]. On the other hand, many studies investigating structure-function relationships find weak to moderate sized correlations in the same upper range that we found (r = .31 to .39) [8,9,70,71,75]. Interestingly, in those studies finding much greater structure-function correspondences, the populations tend to be older [7,76–78] or have a disorder that impacts the brain [79]. These increases in correlation strength are often attributed to the greater variability in both structure and function that is necessary to detect individual differences in those populations.

Different sources of noise for both DTI and fMRI also can contribute to the low correlations. Interestingly, Honey et al. [75] compared structure-function relationships using low-resolution maps and high-resolution maps and found that brain image resolution impacted those relationships. Specifically, the strength of the relationship was about half the size in the high-resolution map compared with the low-resolution map (r = .36 vs. .66, respectively). The authors attributed this difference to inter-individual differences in anatomical and functional locations, which benefitted from the “blurring” of the low-resolution maps analogous to “smoothing” of data, which is a common preprocessing step in most neuroimaging analysis pipelines. Consistent with these interpretations, we utilized high-resolution data from HCP, which may have ironically decreased our ability to find strong connections. In balance, we interpret the weak to moderately sized correlations in the present study as real correlations, but also acknowledge that WM integrity obviously is one of multiple factors that impact network complexity.

### 4.5 Conclusions

We suggest that network complexity provides a novel window into the dynamics of brain functioning that mean activity and functional connectivity do not provide. Using multiscale entropy, the present study reveals new relationships between brain structure and function on multiple time scales. While evidence is gathering that greater WM integrity facilitates information processing in neighboring brain regions, this facilitation may come at a cost or disrupts a balance in how local WM integrity affects information processing across more distributed (distal) brain regions. The present findings support computational simulations of structure-function interactions, but also implicate two separate pathways through which WM integrity affects brain function in the prefrontal cortex—an executive-prefrontal pathway and a perceptuo-occipital pathway. This dual-structural path provides a framework to investigate how lesions or deterioration of these pathways differentially affects cognition in aging or clinical disorders. Future work should be aimed to test how these two pathways impact information processing at the behavioral level both in normal adults as well as changes in behavior as a function of age and disorders.

## Acknowledgments

Data was taken from the Human Connectome Project, WU-Minn Consortium (Principal Investigators: David Van Essen and Kamil Ugurbil). This work was supported by the National Institutes of Health Centers that support the NIH Blueprint for Neuroscience Research [grant numbers 1U54MH091657]; the McDonnell Center for Systems Neuroscience at Washington University. Aspects of these data were presented at the 13^th^ International Conference on Cognitive Neuroscience in San Francisco, CA.

## Author Contributions

IMM conceived and designed the experiments; IMM and JTS analyzed the data; IMM and JTS contributed analysis tools; IMM and JTS wrote the paper.

